# Temporal dynamics and readout latency in perception and iconic memory

**DOI:** 10.1101/2024.10.07.616988

**Authors:** Karla Matić, Issam Tafech, Peter König, John-Dylan Haynes

## Abstract

Following the offset of complex visual stimuli, rich stimulus information remains briefly available to the observer, reflecting a rapidly decaying iconic memory trace. Traditionally, iconic memory decay is assumed to begin with stimulus offset. Instead, here we found that available information begins decaying already when cues are presented in the final stage of stimulus presentation. Using closely spaced (“micro-timed”) readout cues and a theoretical model of information availability, we observed that a cue has to be presented around 10-30 milliseconds before stimulus offset in order to access the full sensory information. We suggest that this does not reflect an early loss in sensory encoding, but instead it is a consequence of a latency in the processing of the cue which postpones the readout of the sensory representation by 10-30 milliseconds. Our analysis also shows that spatial proximity of items in complex arrays impacts sensory representation during both perceptual encoding and initial memory decay. Overall, these results provide a theoretical and empirical characterization of the readout from visual representations, and offer a detailed insight into the transition from perception into iconic memory.

## Introduction

Visual perception has the ability to temporarily retain sensory information even after the removal of a stimulus. A classical finding is that observers presented with large stimulus arrays can only remember around 4-5 items, consistent with the capacity limits of short-term memory (Cowan, 2001). However, if they are prompted very briefly after the removal of the stimulus to report a subset of the presented items (a so-called “partial report”), observers can perform with high accuracy (Coltheart, 1980; Lamme, 2010; Pratte, 2018; Sperling, 1960). This finding suggests that rich stimulus information remains available for report for a brief period even after stimulus offset, stored in a short-lived but large-capacity memory store known as iconic memory (Neisser, 1967).

The sensory trace of stimulus information is suggested to decay exponentially over the course of 300-500 ms following stimulus offset (Bradley & Pearson, 2012; Graziano & Sigman, 2008; Lu et al., 2005). During this time, the information in the sensory buffer can be selectively accessed and transferred into working memory for further processing (Gegenfurtner & Sperling, 1993). The neurocognitive mechanism involved in iconic memory readout is suggested to rely on the same perceptual processes as the ones used for reading out of “online” physical stimulus representations (Barban et al., 2013; Loftus et al., 1985; Loftus & Hogden, 1988; Teeuwen et al., 2021). In many regards, the iconic representation can be considered a fading continuation of the perceptual (i.e., physical stimulus) representation.

The decay of stimulus information is often assumed to begin with stimulus offset, starting as soon as the stimulus is removed from the screen. However, several studies report that performance is lower when cues are shown at the offset of the stimulus compared to the cues shown before or simultaneously with the stimulus (e.g., Averbach & Coriell, 1961; Bradley & Pearson, 2012; Lu et al., 2005; Sperling, 1960). This would seem to suggest that the decay of iconic memory is, paradoxically, beginning while the stimulus is still presented on the screen. However, there is a simpler explanation (Figure 1). First, there are time delays in neural processing between the events happening on the retina and in visual cortex. Thus, there is also a time delay between the offset of the target stimulus on the screen and the end of the sensory-driven response (akin to the time delay between stimulus onset and response onset in the visual cortex; Figure 1A). Second, processing of the readout-cue is not instantaneous but will also involve latency (Averbach & Sperling, 1961), which is why a cue presented directly at stimulus offset does not immediately trigger read-out from the sensory representation (Figure 1C). The additional sensory and cognitive requirements involved in the processing of the cue might mean that the cue would need to be presented prior to stimulus offset in order to be fully effective. In this view, a cue presented directly at stimulus offset would initiate the readout only once the stimulus-related information has started to decay (Figure 1D). We will refer to this readout-related processing latency as *cue-readout latency*.

**Figure 1.**
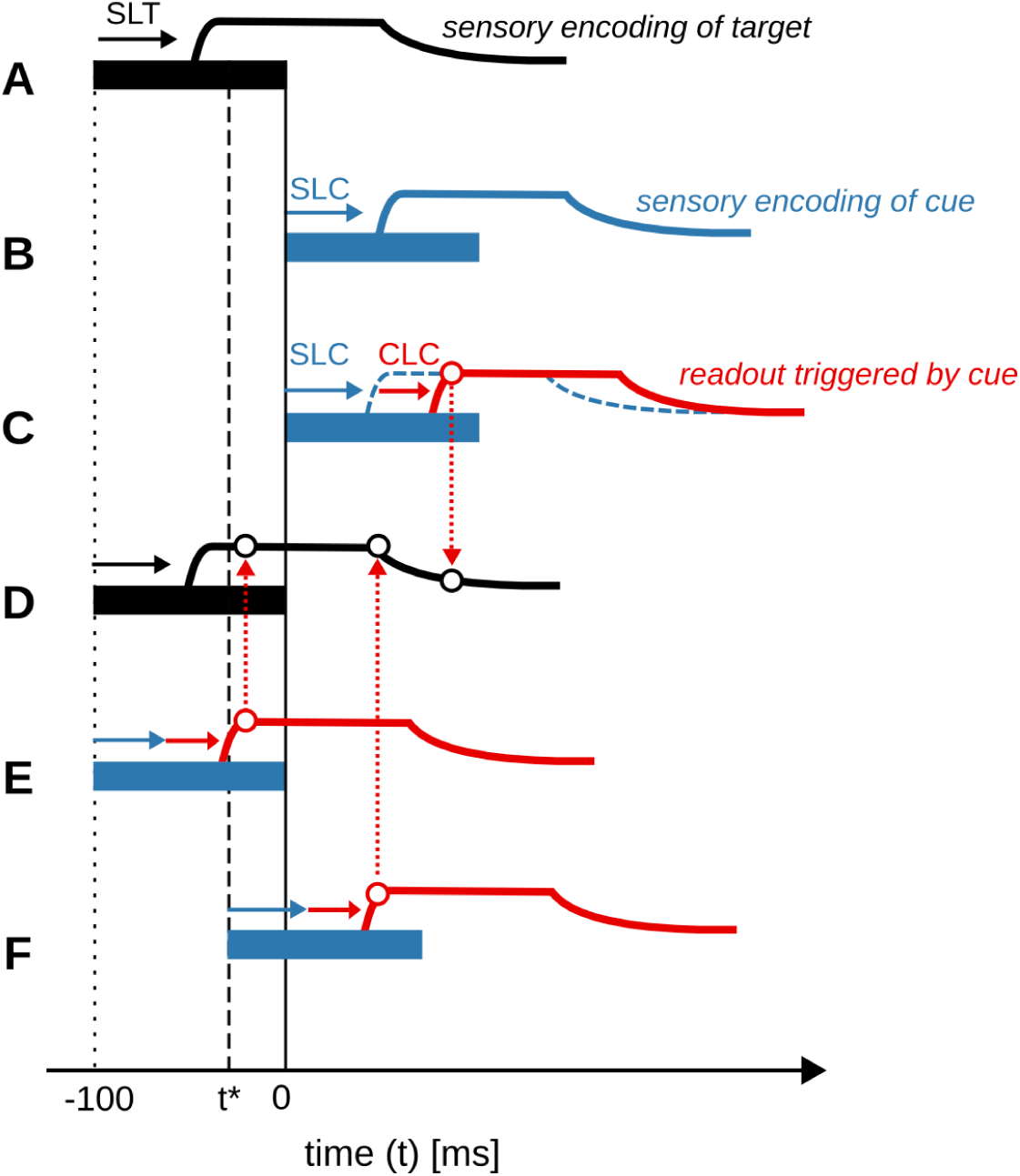
Theoretical neural response latencies. The black rectangle shows a physical target stimulus (e.g. an array of symbols) that is presented on the screen with an onset at t = −100 ms and offset at t = 0 ms. **(A)** After stimulus exposure, stimulus information becomes available in the brain and decays rapidly. However, there is a *sensory processing latency* (SLT) between the target stimulus onset on the screen and the onset of a response in the visual cortex. Thus, the time course of neural information is delayed. **(B)** Here a readout-cue is presented just after the offset of the stimulus. The readout-cue has two relevant processing latencies. The first is the *sensory latency of the cue* (SLC), the latency between the physical onset of the cue and its neural representation in the visual cortex. **(C)** The second processing latency of the cue is the latency between the onset of the cue in the visual cortex and the time it takes to commence the readout from the sensory buffer. We label this the *cognitive latency of the cue* (CLC). **(D)** Readout of sensory information at different latencies (black circles) taps into different levels of information. In the case shown in (C), the readout-cue is so delayed that the readout of sensory information in the visual cortex does not begin before the sensory information about the target has substantially decayed (red circle and rightmost black circle, red dotted line). **(E)** In this case, a readout-cue is presented much earlier, at the same time as the onset of the stimulus. Here, the total cue-readout latency (SLC plus CLC) is short enough for the readout to begin while the stimulus information is still at its maximum. **(F)** Here the readout-cue is presented at a theoretical point in time *t\** such that the readout begins exactly at the latest possible time before stimulus information starts to decay. For this to happen, the cue must be presented prior to offset of the target stimulus with a lead time of (CLC + SLC - SLT). If we simplify by assuming that the sensory latencies of the target (SLT) and the cue (SLC) are equal, then the lead time (i.e., distance between *t\** and stimulus offset) is equal to CLC. Therefore, determining the *t\** would allow us to approximate the CLC.

The aim of the present study was to investigate cue-readout latency in a partial report paradigm with fine-grained temporal resolution. First, in Experiments 1a/1b we systematically replicated the finding that information availability measured already at stimulus offset is lower than the information available during stimulus presentation. Second, in Experiment 2 we further tested the hypothesis that this effect reflects a latency between cue onset and read-out. We expanded the number of manipulated cue timings to “micro-time” the sensory readout, and fit a model specifying the transition from perception into iconic memory including a free parameter estimating the onset of the decay of available information (*t\**). This allowed us to measure the availability of information on the transition between perception and iconic memory with high temporal resolution, and uncover the minimal cue-processing latency required to read information out of a stimulus representation in partial report paradigms.

## Methods

### Participants

We conducted 3 experiments with a total of 82 participants. Participants, aged 18-45 years, were recruited from Humboldt University Berlin’s recruitment system.There were 30 participants in Experiment 1a (17 female), 28 in Experiment 1b (18 female), and 24 in Experiment 2 (19 female). Experiments 1a and 1b differed only with regards to the cue-delay conditions (long delays in Experiment 1a and short delays in Experiment 1b; see Procedure). All participants were untrained, sampled from a diverse pool, and reported normal or corrected-to-normal vision. The study was approved by the Ethics Committee of the Institute of Psychology at the Humboldt University Berlin, and participants were remunerated for their participation with 10€ per hour.

### Visual stimulus and task

The experimental protocol was programmed in MATLAB (version 9.6.0 R2019a; The MathWorks Inc., 2019) using the Psychophysics Toolbox V3 (Brainard, 1997), and ran on a Linux Ubuntu 22.04 computer. Experiments took place in a soundproof experimental booth under dimmed light. Experiments 1a/1b used a 27” LCD monitor with a 1920×1080 resolution and a 60Hz refresh rate for stimulus presentation. In Experiment 2, the stimuli were presented on a high-speed LCD monitor, model LG27GN800, with a resolution of 2560 x 1440 pixels and at refresh rate of 144Hz. The timing reliability of the screen refreshes was monitored internally during the experiments and confirmed using a high-speed camera. Participants were seated 90 cm away from the screen, with their head fixated on a chin rest. Responses were collected on a computer keyboard using the left/right arrows and the return key.

Stimulus arrays were generated by randomly sampling the 21 consonants of the English alphabet without repetition. Letters were shown in a sans serif uppercase font with a size of 0.9 degrees of visual angle. The letters were arranged on a circle around the fixation point at an eccentricity of 4 degrees of visual angle (Figure 2A). In Experiments 1a/1b, the arrays consisted of 4, 8, 12 or 16 letters. In Experiment 2, the array always consisted of 12 letters. The stimulus array was presented for 100 ms in Experiments 1a/1b, and 104 ms in Experiment 2 (the difference was due to different screen refresh rates). Depending on the condition, the cue was shown either before, during, or after the stimulus array (Figure 2B). The cue was a high-contrast line extending from the fixation dot to an invisible circle with a radius of 3.5 degrees of visual angle, designed to extend to the close proximity of the cued letter without masking the stimulus. In Experiments 1a/1b, the cue was always presented for 100 ms, and in Experiment 2, the cue was presented for 49 ms.

**Figure 2.**
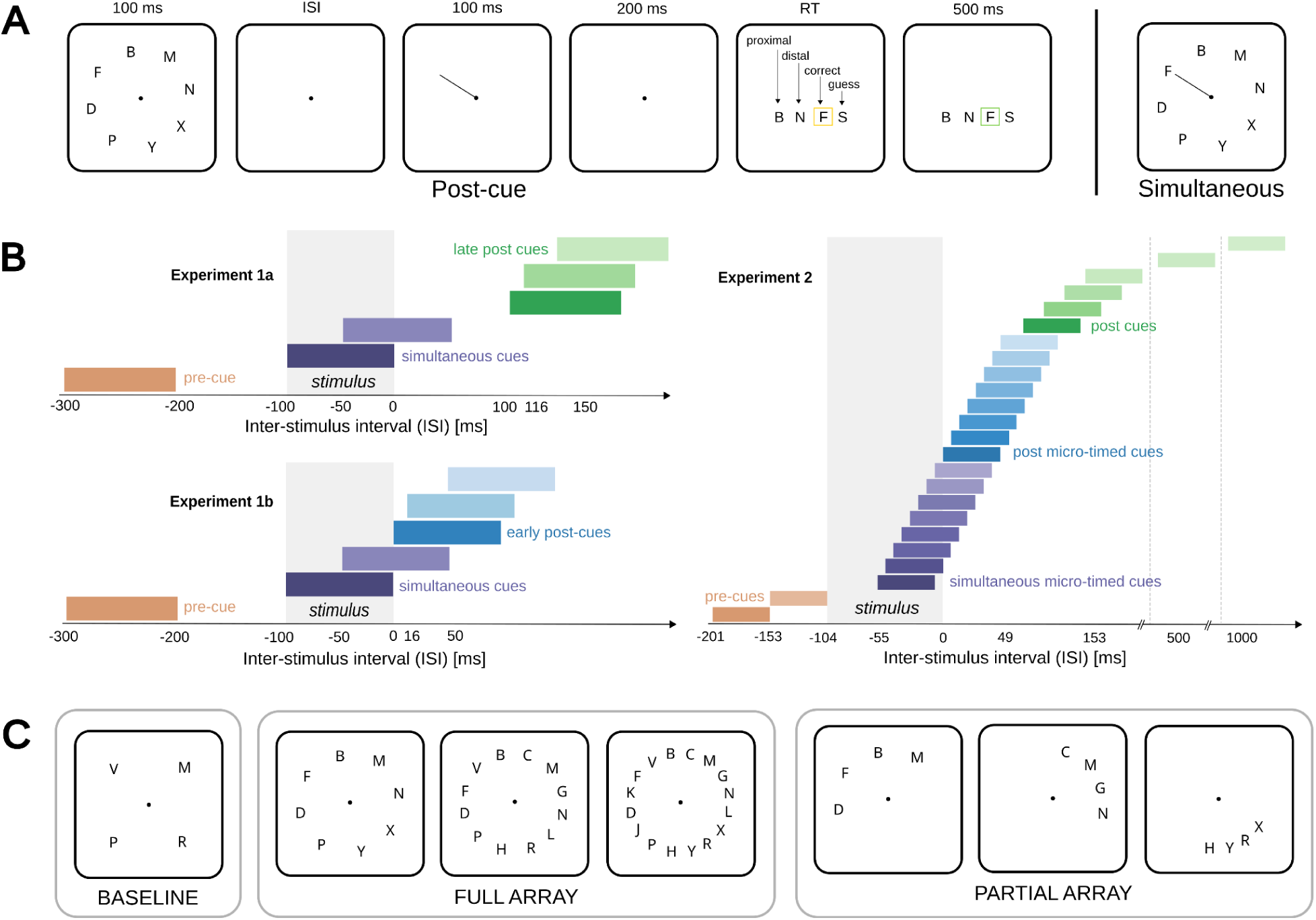
Experimental design, stimuli, and conditions. **A)** All experiments employed a partial report paradigm (Sperling, 1960) with radial target stimulus arrays, a spatial cue, and a response selection screen. Participants first saw a target screen with an array of consonants (for 100ms in Experiments 1a/b, and 104 ms in Experiment 2). On each trial, one of these consonants was cued, which indicated that this was the item to be selected on a subsequent delayed response selection screen. The response selection screen also contained other consonants that were distractors. These were selected to measure substitution errors (adjoining letter to the cued), distant errors (another letter from stimulus array), and guessing errors (letter that was not presented at all). The cue was presented with variable timing relative to the target stimulus arrays (either before, at the same time or after). **B)** Grey rectangles mark the time of target stimulus exposure (timing is relative to target offset at 0ms). Coloured rectangles mark the cue exposure conditions. The distances are drawn approximately but not exactly to scale. In Experiments 1a and 1b (left), cues were presented such that the onset of some cues overlapped with stimulus exposure (simultaneous cues in purple), and in other conditions cues appeared before the stimulus (pre-cue in orange) or after stimulus offset (Experiment 1a - longer ISIs marked in green; Experiment 1b - shorter ISIs marked in blue). In Experiment 2 (right), cues were temporally micro-spaced to sample the readout during and directly after the stimulus exposure with a higher temporal resolution. **C)** In Experiments 1a/1b we manipulated the size of the stimulus array between 4, 8, 12 and 16 letters. In partial array trials, subsets of the full arrays containing only 4 letters at different distances were shown and compared to the baseline condition to examine the encoding and decay of spatial information. In Experiment 2 only the full 12-letter array was used.

Participants were instructed to report the cued letter (i.e. the letter to which the cue was pointing). They indicated their choice by selecting one letter from a response array (Figure 2A). The response array always contained 4 letters: the correct letter; one of the two letters adjoining the correct letter (“proximal” distractor); one of the other letters from the stimulus array, but not adjoining (“distal” distractor); and one letter that was not shown on the given trial (“guessing” distractor). The four letters were shown in a shuffled order, and the initial selection rectangle always appeared at a random location. The response array remained on screen until the participant selected one of the letters. Each trial ended with feedback indicating the correct response with a green rectangle.

### Procedure

In all experiments, the experimental session started with verbal instructions shown on the screen, followed by a practice block of 15 trials. Participants were instructed to focus on the central dot and prioritize accuracy over speed in their responses. Each participant completed 900 trials in Experiments 1a/1b and 1080 trials in Experiment 2, with sessions lasting 60-90 minutes, depending on response speed and break lengths. Between blocks there was an enforced break of at least 60 seconds, and three short breaks of at least 10 seconds equidistantly spaced within each block. Participants were encouraged to take as long breaks as they needed.

#### Experiments 1a/1b

The main experimental part consisted of 6 blocks of 90 trials. Arrays of 8, 12, and 16 array sizes were each repeated twice, and shown in random order. The cue timings are reported with respect to stimulus *offset* (Figure 2B), and in the rest of the text referred to as the inter-stimulus interval (ISI). The stimulus duration in Experiments 1a/1b was 100 ms. Experiment 1a employed six ISIs: pre-cue (−300 ms), simultaneous cues (−100 ms, −50 ms), and late post-cues (100 ms, 116 ms, 150 ms). Experiment 1b employed the same pre-cue (−300 ms) and simultaneous cues (−100 ms, −50 ms), but early rather than late post-cues (0 ms, 16 ms, 50 ms). The manipulation of array sizes was block-wise (i.e., constant within a block), and manipulation of ISIs was trial-wise and counterbalanced across the blocks. The cue timing was therefore unpredictable on each trial. For the purpose of assessing information decay across a larger range of ISIs, the data from Experiments 1a/1b were combined, resulting in a total of 27 experimental conditions (3 array sizes X 9 ISIs). Each subject completed a total of 540 experimental trials, with 30 trials per experimental condition. The complete dataset included a total of 52200 trials, with at least 840 data points per distinct experimental condition.

In addition to the main experimental blocks, each subject also completed 4 blocks with partial arrays (Figure 2C, right). These blocks were inserted in the middle of the experiment (i.e. preceded and followed by 3 main experimental blocks). The purpose of these partial array blocks was to measure the effects of spatial proximity of the letters within an array on sensory readout. To that end, we presented stimulus arrays which only contained 4 letters, and manipulated the proximity of the letters. In the case with no spatial proximity, the letters were shown as far apart as possible. In the other three cases, the 4 letters were shown with the spatial distance mimicking that of the arrays of 8, 12, and 16 items used in main experimental blocks. In simple terms, the 4 presented letters were a “slice” from the larger letter arrays. This spatial distance was constant for a given condition block, and the ISIs were randomized in the same way as in the experimental blocks. Each subject completed 15 partial-array trials per condition and a total of 360 trials. Overall, a total of 28 partial-array conditions (4 spatial distances X 9 cue ISIs) were collected, resulting in at least 420 data points per distinct partial-array condition.

#### Experiment 2

Experiment 2 followed the procedure of Experiment 1 with several alterations. First, only arrays of 12 letters were used. Second, the spacing of the ISIs was micro-manipulated to precisely measure the timing of the onset of informational decay (Figure 2B). To this end, we used a monitor with a refresh rate of 144Hz, due to which the duration of the stimulus exposure was 104 ms (15 frames). The duration of the cue was 49 ms (7 frames). A total of 24 ISIs were measured: 2 pre-cues; 8 simultaneous and 8 post “micro-timed” cues, spaced frame-wise every 6.9 ms between −55 ms and 49 ms relative to stimulus offset; 4 iconic memory cues between 50 and 150 ms, and 2 long cues at 500 and 1000 ms. Third, there was no block-wise manipulation. The only manipulation were the ISIs, which were randomized within and across 12 blocks. Each of the 24 conditions contained 45 trials per participant, resulting in a total of 1080 trials per participant. The complete dataset for Experiment 2 consisted of 22680 experimental trials, with at least 720 trials per condition.

### Data analysis

Data analysis was performed in Python (version 3.12.3). The data and the code are available from the project repository: https://gitlab.com/karlaxmatic/microtiming.

#### Data preprocessing

In Experiment 2, due to a technical error in data collection in 8 out of 24 participants 9 ISI conditions were shown at slightly different times: these cues were micro-timed in the range between 50-150 ms after cue offset. We did not notice any systematic difference between these participants and the rest of the sample. We therefore did not exclude these participants from the analysis.

#### Two-stage model of visual information availability

We presented arrays of letters of varying sizes and asked the observers to report one of the letters selected by the cue in a partial report paradigm. Based on the results of Experiments 1a/1b, we model visual information availability as a two-stage process: a phase that is governed by external stimulus input, and a phase that is based on an internal iconic memory trace. After estimating the shifting point between the two stages (*t\**), in Experiment 2, we empirically validate the model by measuring information availability around the predicted shifting time points. We define the availability of visual information *A* as the average performance *P* multiplied by the size of the presented array *S,* and calculate it for each subject *i,* ISI *t,* and array size *S*:

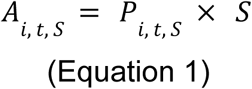

This formulation is equivalent to the calculation of the theoretical capacity of iconic memory (Sligte et al., 2010; Sperling, 1960).

We model the availability of visual information on the transition between perception and iconic memory as a two-stage process: the *exogenous* stage when sensory information is sourced externally from the stimulus, and the *endogenous* stage when information is read out of a rapidly decaying iconic memory. We approximate the first stage reflecting exogenous information availability while the stimulus is still on the screen, termed *a_stim_*, as a constant. We approximate the second, endogenous stage (i.e., the memory-decaying stage) assuming an exponential decay, similarly to previous studies modeling the decay from iconic memory (Graziano & Sigman, 2008; Lu et al., 2005). However, we modify the three-parameter exponential function to incorporate the cue-processing delay. Thus, we allow for the model to reflect a potential drop in the available information already when the cue is presented towards the end of the target stimulus array due to the processing delays of the cue. We model this as an exponential decay that begins already when the cue is presented during the stimulus presentation. To estimate the onset of the decay, we add the fourth free parameter *t\**. The two-stage model of information availability is given as the following piecewise-defined expression:

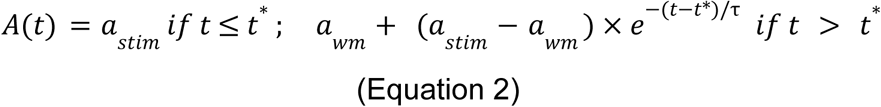

If the cue is shown before a critical time point *t\**, available information will be constant, estimated by the parameter *a_stim_*. At the time point *t\**, available information begins to decay exponentially. Parameter *a_wm_* reflects the asymptote of the memory decay, which can be interpreted as the amount of information transferred to visual working memory (Cowan, 2001). The information available at the onset of the exponential decay (i.e., at *t\**) is equal to *a_stim_*, but only the portion of information that is stored in iconic memory on top of the working memory (*a_wm_*) will decay exponentially. For that reason, the parameter that scales the exponential decay is defined as *a_stim_ - a_wm_*. The parameter *τ* is the time constant of the exponentially decaying information availability, determining the rate of the decay.

The parameter *t\** specifies the onset of the exponential decay. This is the last point in time when a cue can be shown to access the maximal information available from the external stimulus (note that performance is high but not perfect when the cue is presented anytime during the target). If we make a simplifying assumption that the visual processing delays for targets and cues are comparable, then the distance between the time point *t\** and the offset of the stimulus indicates the cognitive component of the cue-readout latency (see Figure 1).

#### Model fit and parameter optimization

We fit the four-parameter model (Equation 2) using a grid-search algorithm minimizing the mean sum of the squares of the residuals (i.e., least-mean-square-error estimation) implemented in Python. The grid search involved looping through a range of values for each parameter, calculating the model-predicted values for all ISIs, and taking the mean squared difference between the predicted and observed values.

In Experiment 1a/1b, we estimate the availability of information during the stimulus and in the first 150 ms following its removal from the screen. We estimated model parameters separately for the arrays of 8, 12, and 16 letters. We specified the following range for each of the parameters: −50 - 0 for *t\**, 1 - 140 for *τ,* and 1 - 7 for *a_wm_*. For *a_stim_*we specified the minimum as array size - 4, and maximum as the array size, so the range was 4-8 for the 8-letter arrays; 8 - 12 for the 12-letter arrays, and 12-16 for the 16-letter arrays. Because there were only 3 post-cue conditions for each subject (either between 100 - 150 ms following stimulus offset in Experiment 1a, or 0 - 50 ms after stimulus offset in Experiment 1b), it was not possible to fit the model individually for each subject. Therefore, we fit the model to the group-averaged availability estimates for each ISI.

In Experiment 2, we followed a closely related procedure. We fit the model to all simultaneous and post-cue conditions. We constrained the parameter grid-space as follows: −50 to −15 for *t\**, 1 - 140 for *τ*, 1 - 7 for *a_wm_*, and 9 - 12 for *a_stim_*. We fit the model to both group-averaged and individual subject data. The individual subject data contains a lot of variability due to individual differences as well as due to measurement noise and relatively few trials per condition. For this reason, we expanded the parameter space to allow for a higher variety of model fits. Specifically, we allowed the parameter *t\** to vary in the range during stimulus presentation (−50 - 0 ms), but also in the range after stimulus offset (0 - 15 ms), to allow for the possibility that some subjects’ individual performance may only begin decaying after stimulus offset.

#### Hypothesis testing

To compare performance in pre-cues and early post-cues with simultaneous cues (Figure 3), we employed two-sided paired-sample t-tests. To test whether parameter *t\** is significantly earlier than stimulus offset in Experiment 2, we tested the mean of parameter estimates of individual subjects (Figure 6) against a ‘population mean’ of 0 ms, representing stimulus offset, using a one-sample t-test. We report significance levels corrected for multiple comparisons using Bonferroni correction. To compare the effects of spatial proximity on information availability, we used ANOVA functions from the Python package ‘statsmodels’. For summary statistics we report mean (M), following standard deviation in brackets. The error bars in all plots indicate confidence intervals.

**Figure 3.**
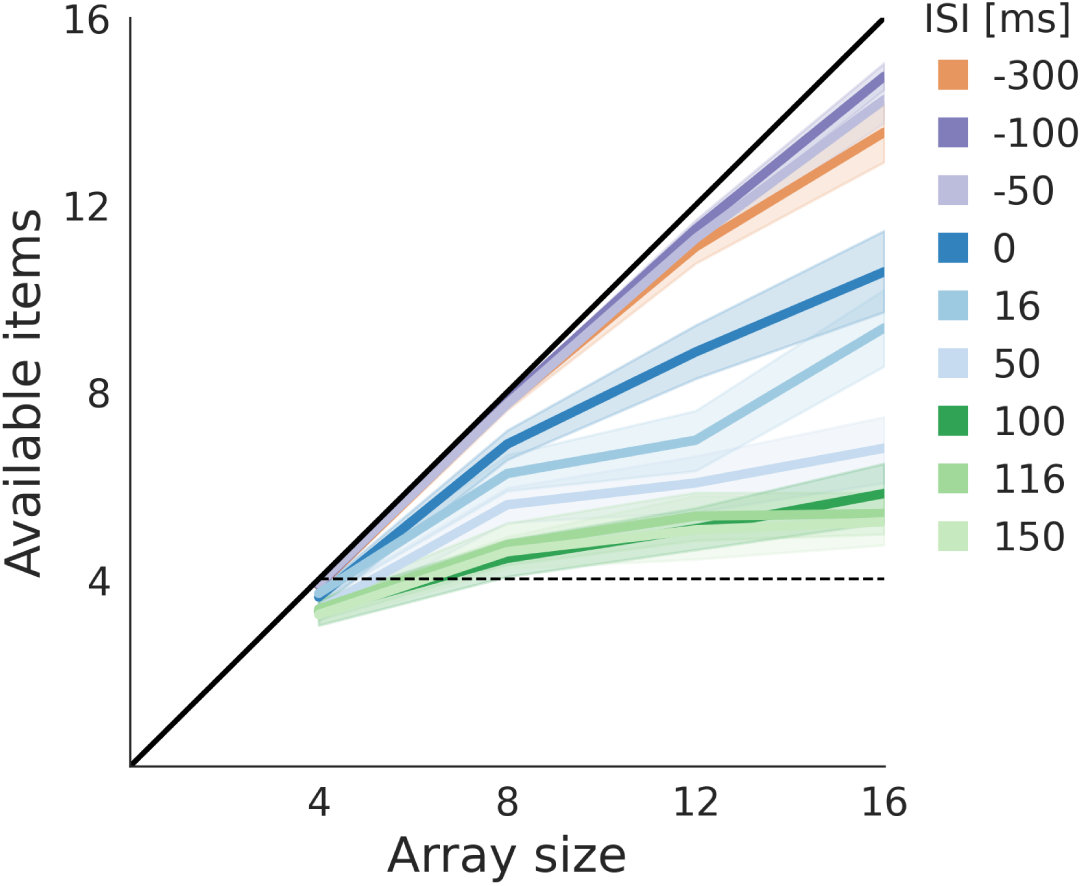
Information availability in Experiments 1a/1b. The plot shows availability of items measured with a pre-cue (orange), simultaneous cues (purple), early post-cues (blue) and late post-cues (green). The availability is plotted relative to the theoretical maximum (black diagonal). When cues were presented before or during the stimulus, performance was close to the theoretical maximum. As predicted, performance was significantly worse in the first post-cue than during the stimulus, and it leveled off around 100 ms. Error bands show 95% confidence intervals.

## Results

### Visual information availability model

In Experiments 1a/1b, participants viewed arrays of 4-16 letters for 100 ms. A single letter was cued for partial report, either before the stimulus (pre-cues), simultaneously (simultaneous cues), or after the stimulus offset (post-cues). Figure 3 illustrates the number of items reported (for a definition see Equation 1) for arrays of 4, 8, 12 and 16 letters, across all measured ISI conditions. Availability was similar for pre-cues and simultaneous cues, approaching the theoretical maximum. However, availability was slightly reduced in the 16-letter array, likely due to visual crowding, which is further discussed in the next section. Notably, availability decreased sharply at the first post-cue. A two-sided t-test revealed higher availability in the last simultaneous cue than in the first post-cue for array sizes of 8 items (t(27) = 5.79), 12 items (t(27) = 9.56) and 16 items(t(27) = 7.99; all p < 0.001, corrected for multiple comparisons). This indicates that availability of visual information suddenly drops sometime between the middle and the end of stimulus presentation.

We hypothesized that the drop in available information at stimulus offset (Figure 3) results from an early onset of the information decay due to an additional latency between the presentation of the cue and the effective readout, which we label cognitive latency of the cue (Figure 1C). We estimated this latency by modeling the information available to the observer as a two-stage process (Figure 4). In the first stage, while the stimulus is shown on the screen, there is a constant amount of information which is supported by direct “online” input from the retina. In the second stage, when the stimulus is removed from the screen, information is sourced internally by reading out of a quickly decaying memory trace of the stimulus (see Methods for details). The early decay presumably reflects an additional latency in cue processing (see Introduction, Figure 1). The two-stage model showed a very good fit to the group-averaged data (MSE = 0.018, 0.028, and 0.039 for 8-, 12- and 16-item arrays, respectively). Figure 4 (table) presents the best-fitting models for the three array-size conditions. The estimated onset of the decay *t\** relative to stimulus offset was 16 ms for 8-letter arrays, 14 ms for 12-letter arrays, and 25 ms for 16-letter arrays.

**Figure 4.**
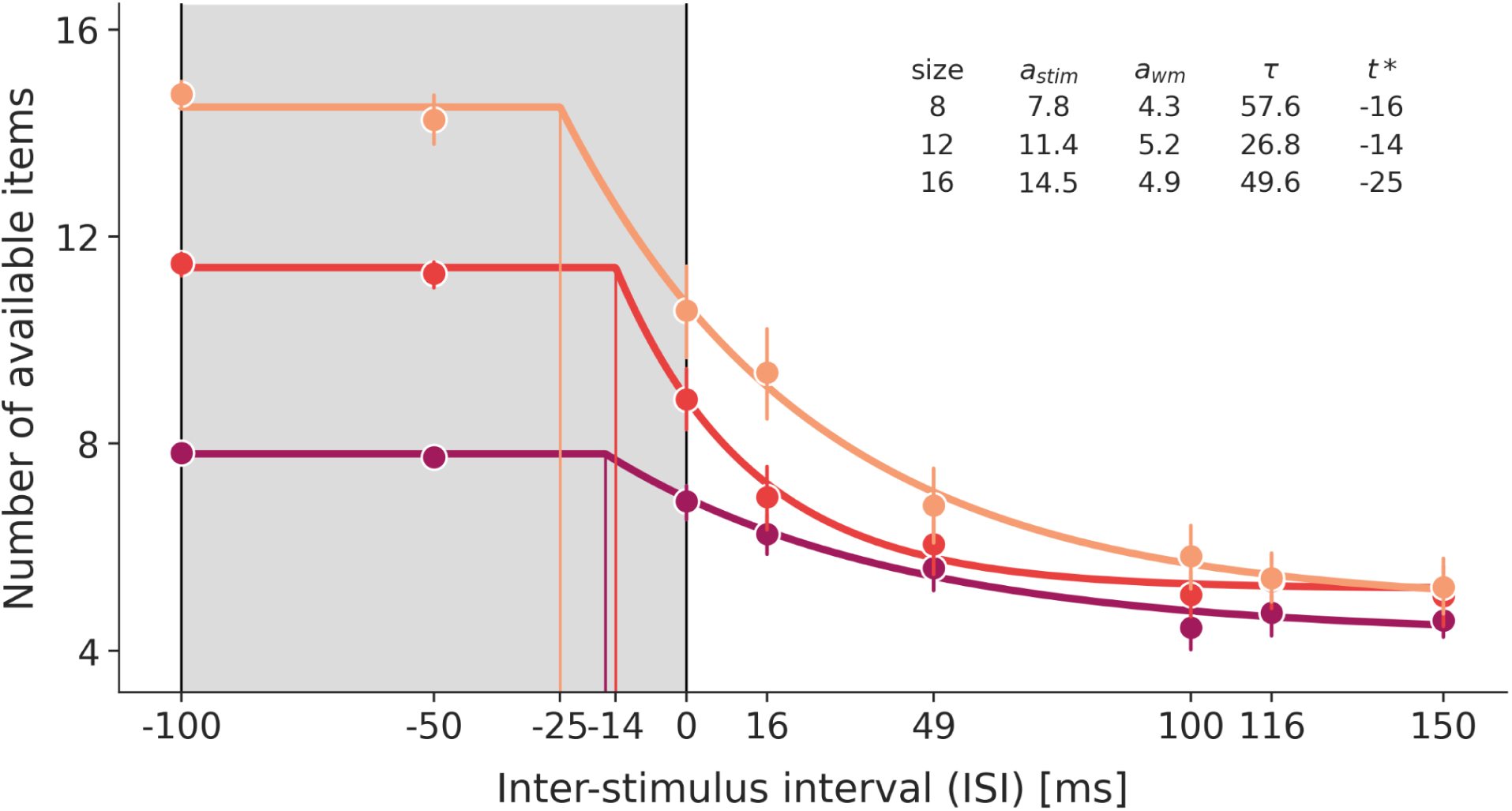
Model of availability of visual information. Theoretical model of information availability was fit to the data of Experiments 1a/1b. The number of available items was measured during stimulus presentation (gray rectangle) and following stimulus removal. Observed information availability was very high during the stimulus presentation (parameter *a_stim_*), but it began to decay before the termination of the stimulus, at time point *(t\**). The estimated *t\** was 16, 14 and 25 ms before stimulus offset for arrays of 8, 12, and 16 letters respectively. The parameter *a_wm_* specifies the asymptote of the model (i.e., the amount of information transferred into visual short-term memory), and the parameter *τ* specifies the rate of exponential decay. Error bars are 95% confidence intervals.

### Micro-timing validates early decay

Results from Experiment 1a/1b (Figure 3) showed significantly lower information availability at stimulus offset compared to during stimulus presentation. The two-stage model’s close fit (Figure 4) suggests information starts decaying before the stimulus offset, likely due to cue processing latencies. However, we could not conclusively validate that information availability indeed decreases before the stimulus offset because we did not measure any data points between the middle and the end of the stimulus. To address this, Experiment 2 replicated early decay and empirically tested the model by capturing measures in the missing 50 ms range before stimulus offset. We maintained the same task and setup, but we used “micro-spaced” cue conditions. With this design, we could directly observe the transition between the external and internal stage of information availability (Figure 5).

**Figure 5.**
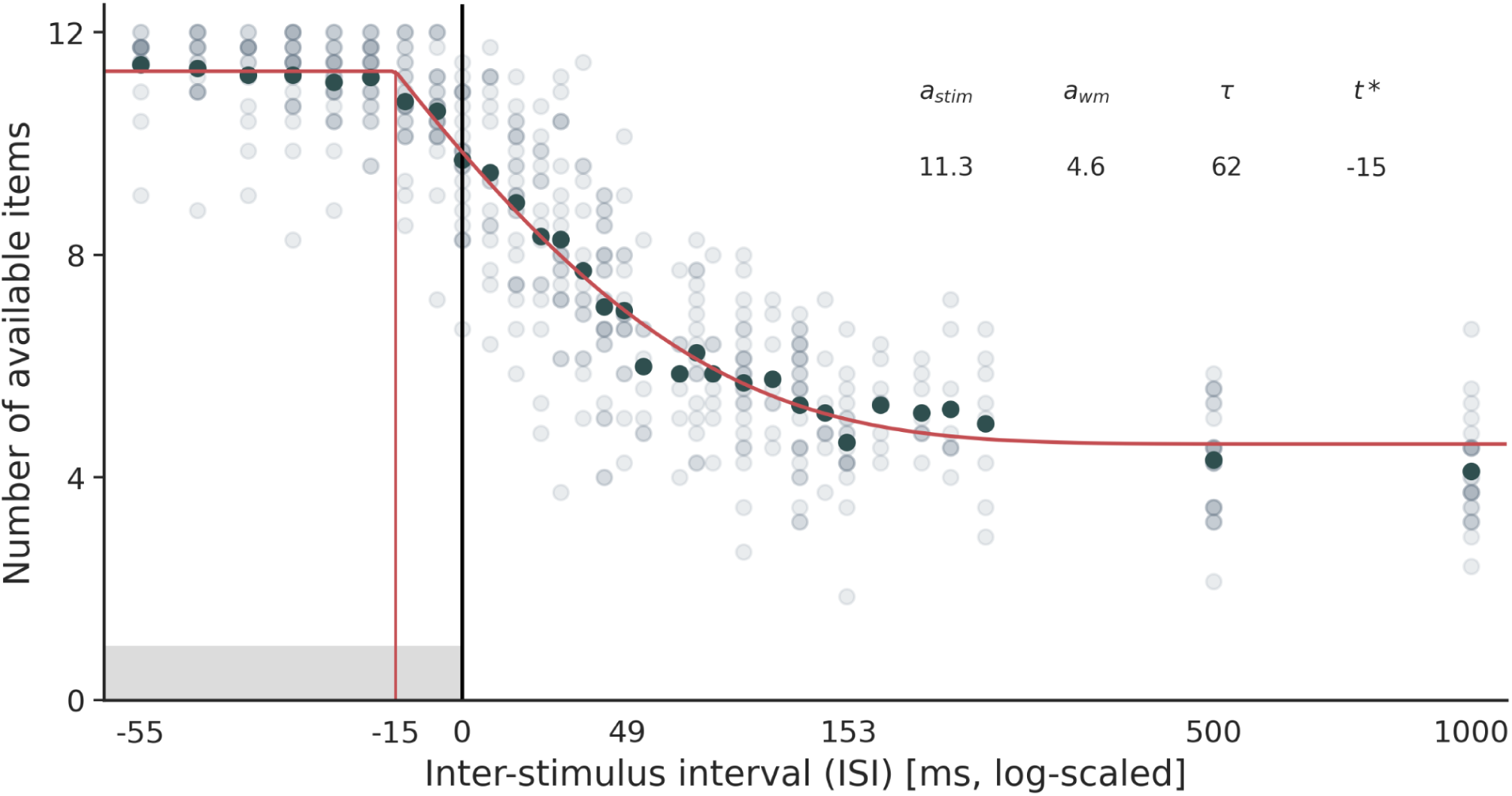
Group-estimated information availability in the micro-timing Experiment 2. We estimated the four-parameter two-stage model of information availability by fitting it to group-averaged performance for each ISI (see Methods). The performance of individual subjects is shown in gray, with the group-averages in darker gray. The model prediction is shown in red, and the four model parameters are given in the table. The shifting point *t\** is estimated at 15 ms before stimulus offset. We explain the distance between the shifting point *t\** and the end of the stimulus as the duration of the cognitive latency of the cue (see Figure 1 for details). The x-axis is converted to a logarithmic scale to improve the visibility of the large range of cueing conditions.

From the onset of the stimulus and through the mid-stimulus cues, availability of visual information remained constant, indicating that the cue was accessing a stable perceptual representation. The model that showed the best fit to the group-averaged data (average squared error of 0.077), shown in red in the Figure 5, had the following parameters: a_stim_ = 11.3, a_wm_ = 4.6, t* = −15, and τ = 61.56. As predicted by the model fit to the data of Experiments 1a/1b, the measured information availability began decaying before the termination of the stimulus, at 15 ms before stimulus offset (compared to 14 ms in Experiment 1a/1b for the comparable condition with 12-item target displays).

We further fit the same model to the data of individual participants (Figure 6A). Despite each participant having only participated in one data collection session, we observed very good model fits even for this noisy data (MSE ranging between 0.136 - 0.738). The model parameters of individual subject fits are shown in Figure 6B. The onset of the decay (t*) was variable across participants, but was on average significantly lower than 0 ms (i.e., offset of the stimulus; t(23) = −4.24, p = 0.0015). Some participants exhibited maximal information availability up to and slightly beyond the stimulus offset, suggesting a ‘negative’ readout latency where decay onset follows stimulus offset. This may indicate random variability, or that these participants managed to extend the stimulus representation beyond its physical duration, allowing the cue to access maximum information despite readout latency. Training or prior exposure to rapid visual stimuli could influence both stimulus representation duration and processing latencies, as illustrated in Figure 1.

**Figure 6.**
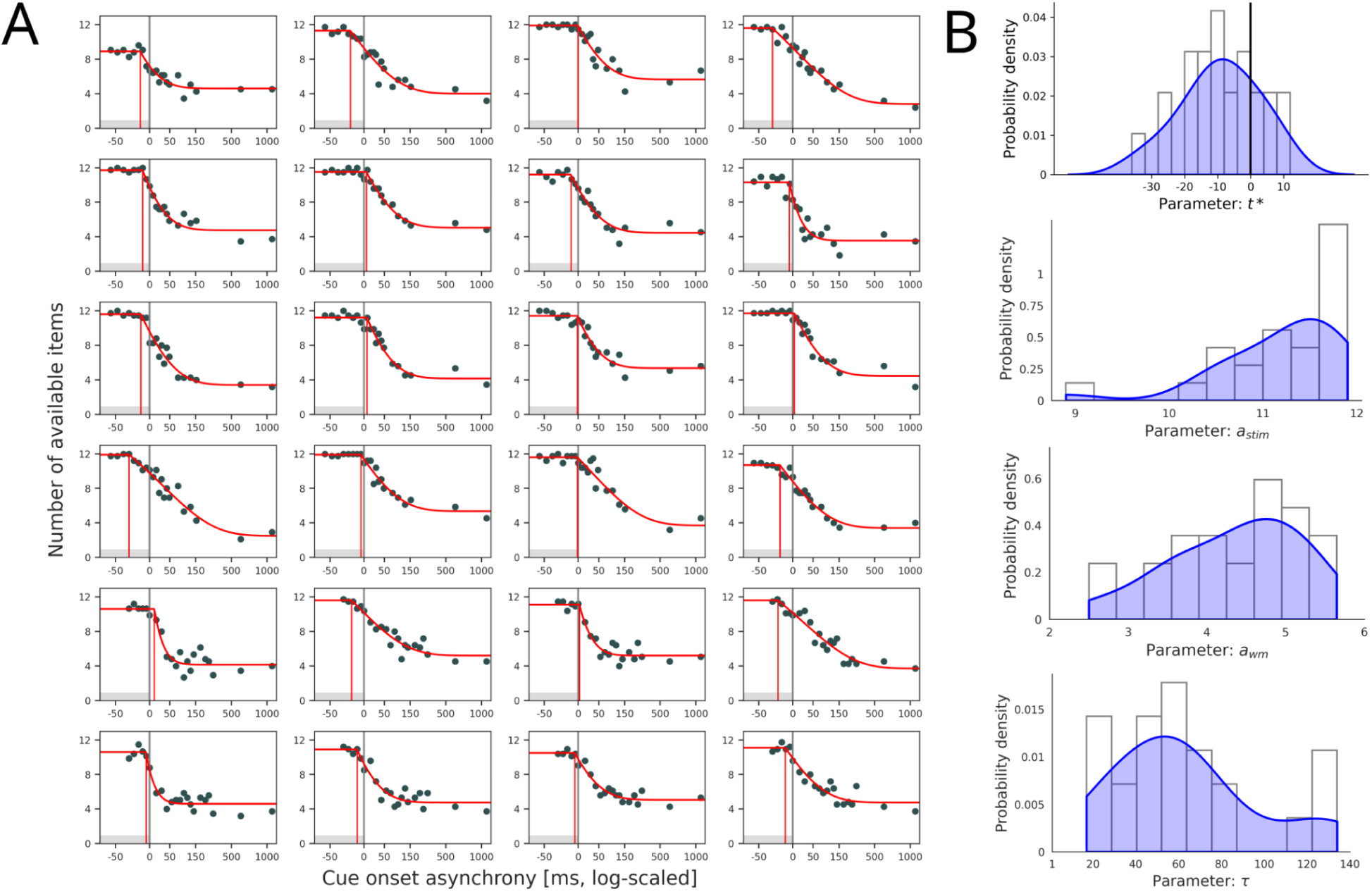
Information availability estimated for individual subjects in the microtiming experiment. We fit the two-stage model of information availability to the individual subjects’ performance in the micro-timing experiment (Experiment 2). **A)** The observed data (in gray) and the model fits (in red) for each of the 24 subjects. The vertical red line indicates the shifting point *t\** estimated for each subject. The x-axis is converted to a logarithmic scale for plotting to improve the visibility of the large range of cueing conditions. Eight participants (in the bottom rows of the plot) saw cue conditions at slightly different times, which is why there are less simultaneous-cues and more post-cues in those participants (see Methods for details). This did not seem to affect the parameter estimates. **B)** Distributions of individual subject estimates of the four model parameters: *t\** (top plot), *a_stim_* (middle-top plot), *a_wm_* (middle-bottom plot, and *τ* (bottom plot). For some participants, the shifting point *t\** was estimated to be after stimulus offset (black vertical line in the top plot), but on average, the shifting point *t\** was estimated to be significantly earlier than the stimulus offset (t(23) = −4.24, p = 0.0015).

### Visual information availability and crowding

We then proceeded to distinguish between several aspects that might contribute to the loss of sensory information over time. Above, we noted that partial report task performance depends on both ISI and array size, even when readout begins during stimulus presentation. A univariate repeated-measures ANOVA of the performance when the cue is simultaneous with stimulus onset (ISI= −100) showed a significant effect of array size (F(3,171) = 8.99, p < 0.001), suggesting that that the number of items in the stimulus array plays a role not only in how much information is retained in memory, but also in how much information can be accessed while the stimulus is still present on the screen. However, a confounding factor is that item proximity increases with more items shown, up to our 16-item maximum. Thus, accuracy loss could stem from either increased item numbers (memory load) or reduced spatial distance (visual crowding).

To assess the effects of visual crowding (Whitney & Levi, 2011), our design employed a separate condition in Experiments 1a/1b with partial-arrays (see Figure 2 and Methods for details). These partial-arrays always showed 4 letters in the stimulus array, thus keeping the total memory load constant, but the distance between the items was manipulated to mimic that for arrays of 8, 12, and 16 letters (Figure 2C). In this way, we could separate the effect of spatial proximity from the effect of memory load. We found that increasing spatial proximity was accompanied by lower performance in the partial report task (main effect of spatial proximity, F(3,1128) = 46.58, p < 0.001), even though the stimulus arrays always consisted of 4 letters (Figure 7A). The accuracy was also decreasing over time (main effect of cue delay, F(7,1128) = 96.1, p < 0.001). Importantly, there was a significant interaction between spatial proximity and cue delay (F(21,1128) = 2.29, p < 0.001), suggesting that information in more crowded arrays decays at a different rate than in less crowded arrays.

**Figure 7.**
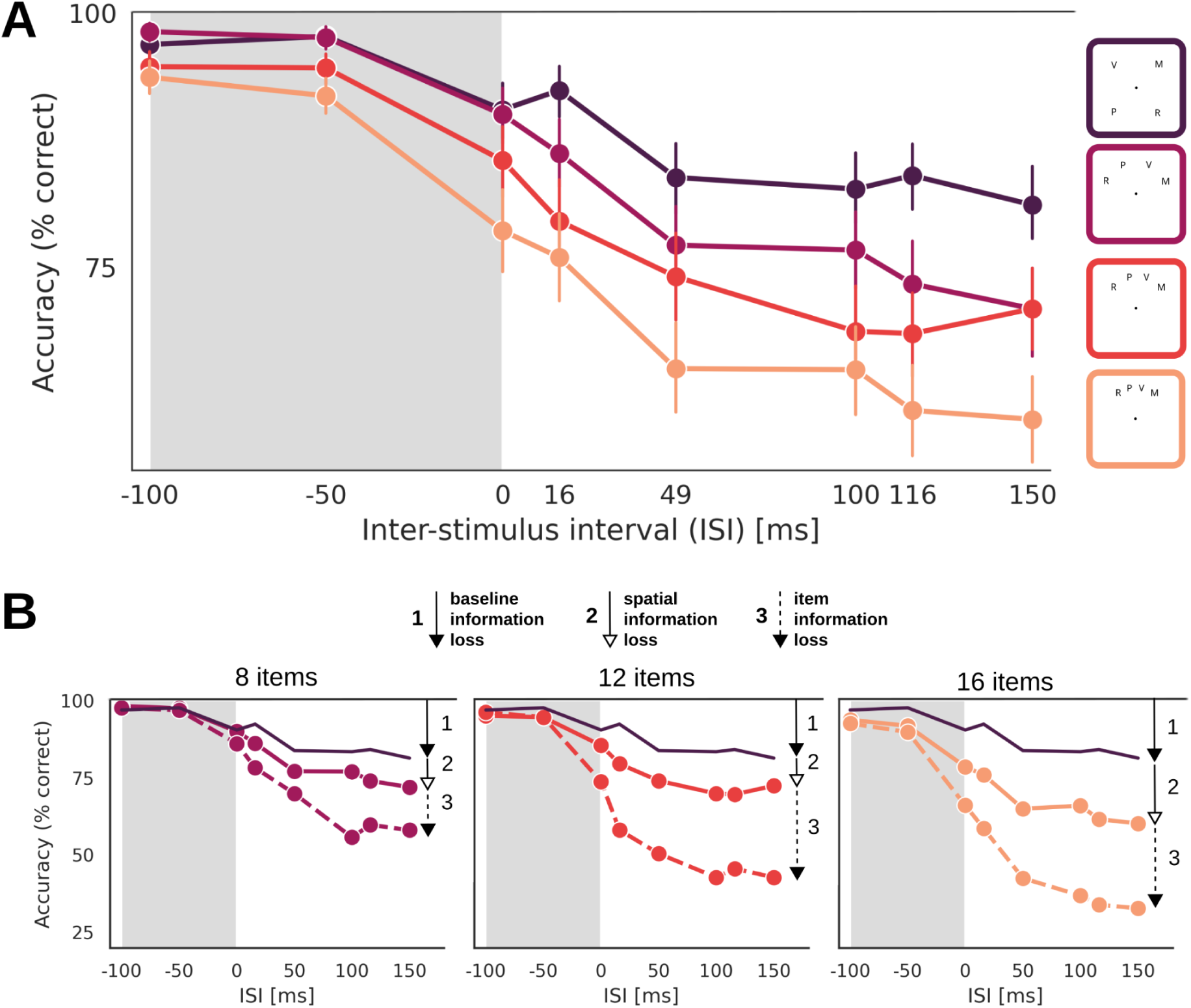
Contributions of different causes to information loss over time. **(A)** In Experiment 1a/1b, we also measured performance for four separate ‘partial-array’ conditions, which all comprised 4 letters, but were arranged in a way that their spatial proximity matched the arrays of 4, 8, 12 and 16 items (color-coded here as purple, pink, red and yellow). Accuracy of the partial report decays faster when items are closer together, suggesting that spatial proximity might also contribute to differences in the displays with full arrays (Figure 3). Error bars show 95% confidence intervals. **(B)** Here we replot the findings from above to reveal any performance differences between the partial-arrays and the matched full-arrays. The three panels show performance for arrays of 8, 12 and 16 items (from left to right). The dark solid lines are the same in each plot and show the 4-item result from panel A for reference. The performance drop compared to online perception (“1”) is a reference for a baseline drop in performance for a non-crowded 4-element display. The plots also show performance in the respective partial-array conditions replotted from panel A (full lines). This reveals the additional drop in performance (“2”) due to spatial proximity (each of these curves used the same total number of 4 items). Finally, the dashed line shows performance in the full array conditions. This reveals an additional drop in performance related to increasing the number of elements even when spatial proximity is kept constant (“3”). Please note that this figure plots accuracy instead of availability to allow direct comparison between full and partial arrays.

We then divided the total information loss into three contributing factors: (a) a baseline information loss (e.g., due to attentional lapses or similar), (b) a loss due to increased spatial proximity (crowding), and (c) a loss due to increasing number of elements. The baseline was defined as the performance for the widest-spaced partial array in which the 4 letters are shown in the diagonal corners of the visual array (Figure 7A, right top; 7B, arrow number “1”). The loss due to increased spatial proximity is defined as the difference in performance between the baseline and the increased crowding conditions with a constant load of 4 items (Figure 7B, arrow number “2”). The loss due to increased item load is defined as the difference in performance between full arrays and the corresponding partial array (Figure 7B, arrow number “3”). It is apparent that all of these factors contribute to the loss of information across time.

Finally, we analyzed error distribution. Considering spatial proximity effects (Figure 7B), we examined its impact on proximal substitution errors, where an adjacent letter is mistaken for the cued letter. A higher proportion of these compared to other errors would suggest that the participant gave a mistaken response because the precise spatial location of the cued item was not available. Figure 8 shows the proportion of three error types: proximal errors (when adjacent-to-the-cued letter is selected), distal error (when a non-adjacent letter is selected), and guessing (when a letter that was not in the stimulus array is selected). In Experiments 1a/1b, the proximal substitution errors were much more prevalent than distal errors and guesses, for all array sizes and with an increasing effect for longer cue ISIs. There was no difference between distal errors and guesses, suggesting that there was no intrusion by the distal distractors on the choice and that any choice of a distal distractor reflected pure guessing. The same pattern was observed in Experiment 2. The more fine-grained time-resolution illustrated that the increase of the proximal errors starts already during stimulus exposure, and remains higher even for long cue delays.

**Figure 8.**
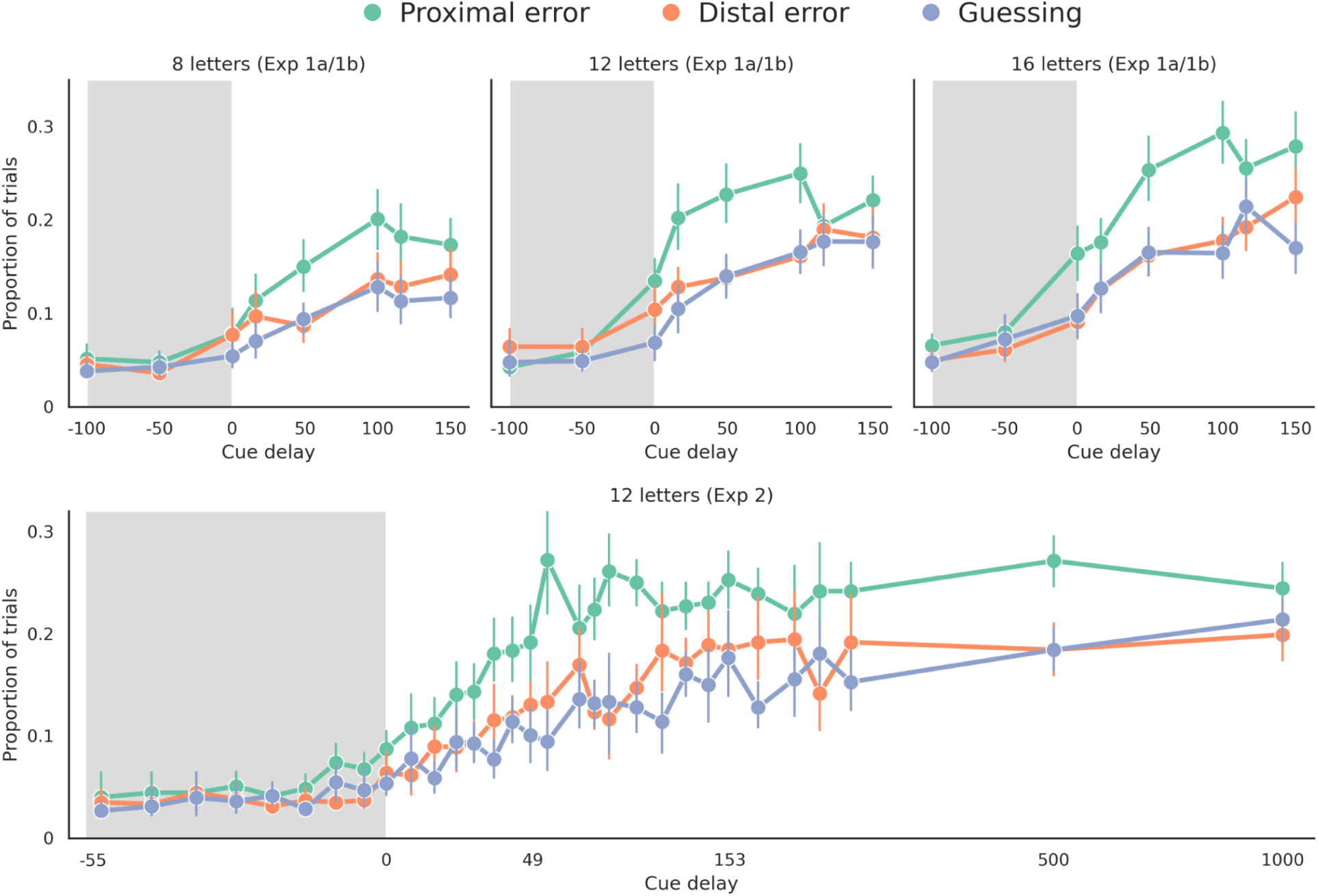
Analysis of error types. Our task required participants to pick the letter indicated by the cue by selecting the correct answer from a response array that contained the correct letter together with three different types of distractors (see Figure 2 for design details). The distractors were: (a) the letter adjoining the cued letter, (b) another letter from the array but not adjoining, and (c) a letter that was not shown on the given trial. We label selection of the neighboring letter a “proximal error”, selection of another letter from the array a “Distal error”, and selection of a new letter that was not shown as “guessing error”. **Top:** The plots show the proportion of error trials in Experiments 1a/1b for arrays of 8, 12 and 16 items. The total proportion of error trials rises over time, proportionally to the decay of information availability. The largest proportion of error trials is proximal errors, and this proportion is higher in larger arrays, and it increases over time. On the other hand, distal errors and guessing constitute a similar proportion throughout and rise together. Thus, participants do not distinguish between novel and distal elements, suggesting there is no intrusion by distal elements, but a substantial intrusion by the neighboring element. **Bottom**: The plot shows the proportion of error trials in Experiment 2, thus replicating the result from the 12-letter condition in Experiments 1a/1b shown in the middle top panel. Micro-timing illustrates that the proportion of proximal error begins increasing when readout is cued already during stimulus presentation. This may suggest that the fine-grained spatial information is the first to begin decaying following the removal of the stimulus from the screen. The x-axis is converted to a logarithmic scale for plotting to improve the visibility of the large range of cueing conditions.

## Discussion

Following the offset of a stimulus, sensory information remains briefly available to the observer (Coltheart, 1980; Sperling, 1960). We investigated when this available information begins to decay relative to the presentation and offset of a stimulus. By presenting a readout-cue in time steps around the offset of stimulus arrays of different sizes in Experiments 1a/1b and further refining the measurement intervals (“micro-timing”) in Experiment 2, we observed that available information begins decaying already when cues are presented between 14 and 25 milliseconds before stimulus offset (Figure 4, Figure 5). We suggest that this puzzling effect might be a consequence of a processing latency between the presentation of the cue and the informational readout from sensory memory.

### Early drop in information and cue-readout latency

Figure 1 illustrates the different latencies involved in the processing of a target stimulus array. The first latency, sensory latency of the target (SLT), results from the time delay between the signals on the retina and visual cortex, which is on the order of around 80 ms in the human visual cortex (Martin et al., 2019). A cue of the same contrast as the target should be registered in the early visual cortex with a comparable latency (sensory latency of the cue, SLC). However, once the cue is encoded in the visual cortex, it takes additional time before readout can begin at the position indicated by the cue. We refer to that additional time as the cognitive latency of the cue (CLC). Thus, total cue-readout latency—the delay from cue presentation to information readout—includes both sensory (SLC) and cognitive (CLC) components. If we assume for simplicity that the stimulus and the cue take a similar amount of time to reach the visual cortex after being registered on the retina (i.e., that SLT = SLC), then the onset of the behaviorally measured informational decay may be taken to reflect the duration of the CLC (see Figure 1F).

In Experiments 1a/1b, we measured information availability for stimulus arrays of 8, 12 and 16 elements (Figure 2) and then fit a four-parameter model of information availability (Equation 2). The model showed a close fit to data (Figure 5, Figure 6), and the parameters of the model are readily interpretable. The parameter a_stim_ provides an estimate of how much information was perceptually encoded from the stimulus, and it suggested that not all elements in larger arrays were available to the observer even during stimulus presentation, potentially due to visual crowding (Whitney & Levi, 2011). The parameter a_wm_ was similar for all three array sizes, suggesting that 4-5 elements get transferred into short-term memory independently of how many elements were initially presented, in line with previous work (Cowan, 2001; Gegenfurtner & Sperling, 1993; Lamme, 2010; Sperling, 1960). Last, the parameter t* provides an estimate of the time when a cue must be presented in order to start reading out information from visual cortex while the stimulus representation has not yet begun decaying, as discussed above and in Figure 1. More specifically, it is an approximation of the duration of the cognitive component of the readout latency (CLC, see above). We observed this to be on the order of 10-30 ms. The exact latencies were varying depending on array size (16 ms for 8-items, 14 ms for 12-items and 25 ms for 16-items; Figure 4), suggesting that it may take slightly longer time to read information out of larger and more crowded stimulus arrays.

In Experiment 2, we finely spaced (“micro-timed”) the readout cues to measure information availability with a finer temporal resolution both during and after stimulus exposure (Figure 2B, Figure 5). Fitting the same model, we found similar best-fitting parameter estimates as in Experiment 1a/1b. Notably, the estimated cognitive latency was 15 ms, thus closely reproducing the estimates of Experiment 1a/1b (where the cognitive latency was 14 ms for the comparable 12-item target display).

The data presented here suggest a short cognitive component of the cue-readout latency on the order of 10-30 ms. This latency is noticeably shorter than the latencies ascribed to the shifts of visual attention, which have been estimated on the order of 100-300 ms (e.g., Reeves & Sperling, 1986; Logan, 2005 Carlson et al., 2006). The large range of these estimates illustrates the fact that the exact experimental setup plays a significant role in determining the estimates, and our paradigm differs from the attentional speed paradigms in several regards. Our experiments employ post-cues rather than pre-cues, and thus the attentional shift may be more efficient. In addition, our estimates are not based on reaction times, as we are estimating the readout latency indirectly from information availability. Last, our paradigm does not require a suppression of a previously attended target, which may further increase the speed of the attentional shift. Given the large disparity between our post-cue partial report paradigm and the designs classically employed to study the speed of attentional shifts, the estimates we report here are likely not directly comparable.

Looking at the observed early information decay, one alternative interpretation may be that sensory components of stimulus and cue processing are not the same, and that the cue simply takes 10-30 ms longer than the stimulus to reach the visual cortex (i.e., that SLC is longer than SLT). However, the cue is more foveal, larger and less complex than the target stimuli. If there were a difference between the sensory components of the cue and stimulus processing, it would be more likely that the stimulus is registered with a longer latency than the cue, rather than the other way around. Yet another alternative explanation may be that some participants are performing poorly overall, since they are not extensively trained, and are thus dragging down the information availability estimates, resulting in what seems like an early information decay. However, the model fits to individual participants’ data (Figure 6) clearly suggest that this is not the case, as the majority of participants exhibit early informational decay.

### Visual information decay and crowding

Another important question was whether available visual information is affected by visual crowding, especially in larger target arrays. Visual crowding is known to affect both perceptual encoding (Townsend et al., 1971; Whitney & Levi, 2011) and visual working memory representations (Tamber-Rosenau et al., 2015), but it might also affect iconic memory. Here we investigated the differences in visual information availability at different levels of crowding during online perception and in iconic memory.

We found that spatial proximity between adjacent letters affects information in iconic memory even when memory load is controlled for (Figure 7). In a separate condition with partial-arrays, we showed a “slice” of stimulus arrays of 8, 12 and 16 letters such that the slice always contained only 4 items (see Figure 2C). This condition thus manipulated crowding independently of memory load. Looking at Figure 7A, we see that there are two conditions (the top two) that have the same amount of information encoded during the perceptual stage, but where information in the more crowded condition decays more quickly. This suggests that the faster drop in iconic memory for crowded displays is not merely a reflection of a lower availability of perceptual information. In addition, comparing performance between partial-arrays and baseline (Figure 7B), we see that spatial proximity between items affects information availability to a similar extent in small and medium stimulus arrays (8 and 12 items), but much more strongly in 16-item arrays. These observations suggest that crowding may contribute to the decay of information during iconic memory, potentially in a process that is different from crowding during perceptual encoding.

While visual crowding in perceptual encoding is related to processes such as lateral interaction or surround suppression between nearby items (Townsend et al., 1971; Whitney & Levi, 2011), crowding in iconic memory may additionally involve increased demands in directing attention to a location. It may be easier to guide attention to resolve individual stimuli when they are still physically present than when attention has to be directed to select the location of a memorized item. In other words, information could still be encoded in iconic memory, but less accessible for readout.

This interpretation is further reinforced by the analysis of errors (Figure 8). We found that spatially proximal errors (i.e., choosing the letter directly adjacent to the cued letter in the response array) become more probable than other types of errors (i.e., choosing another letter from the array or guessing) over the increasing readout delays. In the micro-timed delays we also saw that this increase begins already around the offset of the stimulus. This suggests that encoded stimulus information grows more inaccessible to the observer not only due to the loss of items from iconic memory, but also due to lateral interactions between stimulus elements.

However, with our data we cannot fully resolve the alternative possibility that the crowding effects during memory decay are inherited from limitations in “online” sensory encoding in crowded displays. The two conditions mentioned above (top two lines in Figure 7A) which show equal performance during sensory stage and differences only during memory decay could also be explained by a ceiling effect, because performance during sensory stage is at maximum. Addressing the question of the effect of visual crowding on sensory representations during encoding and iconic memory would be a fruitful avenue for future research.

## Conclusion

Using a temporally fine-grained analysis of the readout from perceptual and iconic memory representations, we found that information initially available to the observer seemingly begins to decay already before the offset of the stimulus. We explain this as the effect of a readout-related cognitive latency on the order of 10-30 ms which effectively delays the readout cue. We also observe that visual crowding contributes not only to the encoding, but also to the decay of visual information. These findings introduce an estimate of the duration of readout latencies in partial report paradigms, and offer a more precise model of the transition between perception and iconic memory.

## Acknowledgements

K.M. was funded by the Max Planck Society and BMBF (as part of the Max Planck School of Cognition). J.D.H. was supported by the Deutsche Forschungsgemeinschaft (DFG, Exzellenzcluster Science of Intelligence); SFB 940 “Volition and Cognitive Control”; and SFB-TRR 295 “Retuning dynamic motor network disorders using neuromodulation”. We thank Karolis Degutis, Kai Görgen and Leonardo Pettini for their input at various steps of this project. We also thank the participants of the MESEC (Mediterranean Society for Consciousness Science) Workshop 2023 for helpful discussions.

